# Warmer climate threatens the occurrence of giant trees in the Amazon basin

**DOI:** 10.1101/2025.02.22.639664

**Authors:** Robson Borges de Lima, Cinthia Pereira de Oliveira, Diego Armando S. da Silva, Daniela Granato de Souza, Lorraine Aparecida Gonçalves, Caroline da Cruz Vasconcelos, Anderson Pedro B. Batista, João R. de Matos Filho, Joselane P. Gomes da Silva, Iran Jorge Correa Lopes, Jadson Coelho de Abreu, Perseu da Silva Aparício, Carla S. Campelo de Souza, Jean Pierre Ometto, Eric Bastos Görgens

## Abstract

Giant trees in the Amazon play crucial ecological roles by acting as substantial carbon sinks and supporting diverse forest ecosystems. However, these emergent trees are increasingly vulnerable to climatic changes, and recent research suggests that the distribution and niche of many emergent species are severely threatened. This study employs ecological niche modeling using data LiDAR and forest inventory, global repository data, and bioclimatic variables to project the potential larger tree distributions under past, current, and future climate scenarios (SSP1-2.6 and SSP5-8.5) in different biogeographic provinces in the Amazon basin. We used algorithms MARS, RandomForest, MaxEnt, and GAM to assess the impact of critical climatic factors, including isothermality, maximum temperatures, and precipitation patterns, on the habitat suitability of these larger trees and species *Dinizia excelsa* and *Goupia glabra*. Our results show that the tall trees dataset, *Dinizia excelsa* and *Goupia glabra* exhibit congruent distinct responses to climate variables, with *Dinizia excelsa* being more sensitive to increased temperature extremes, particularly in the Guiana Shield and Roraima Provinces. In contrast, *Goupia glabra* displays a broader tolerance to precipitation variability. Under the high-emission scenario (SSP5-8.5), both species are projected to lose up to 45% of their suitable habitats by 2080, primarily in southern Amazon provinces like Xingu-Tapajós, where increased drought frequency and temperature extremes are expected. Conversely, the low-emission scenario (SSP1-2.6) suggests potential habitat stability or even slight expansions in the northern Amazon regions due to moderate temperature and rainfall conditions. These findings highlight the urgent need for conservation strategies to protect critical refugia and enhance ecosystem resilience. By integrating niche modeling with climate projections, our study provides vital insights into managing Amazonian giant trees, emphasizing their potential role in mitigating global climate change impacts.

## 1 Introduction

The big trees in the Amazon, with their superlative characteristics, play an essential role in maintaining the structure and functionality of tropical forests. With their high biomass, they act as significant carbon sinks, both on a global and biome scale, directly contributing to climate change mitigation. These trees also regulate the microclimate and support a complex network of ecological interactions, sheltering and sustaining high biodiversity (Lindenmayer & Laurance, 2016; Lutz et al., 2018; Pinho et al., 2020). However, the increasing pressure of global climate change poses a significant threat throughout the Amazon basin (Flores et al., 2024), particularly requiring a more detailed understanding of how these critical climate transitions impact the distribution and survival of the largest trees.

Although the functional characteristics of many species suggest tolerance to environmental variations and climatic gradients (Esquivel Muelbert et al., 2016; Esquivel-Muelbert et al., 2019; Sanchez-Martinez et al., 2024), recent studies indicate that emergent trees are particularly susceptible to climatic variations, such as changes in isothermality, average temperature of the wettest quarter and precipitation in the driest periods (Gorgens et al., 2021; He et al., 2022; Zuidema et al., 2022). For example, these trees, with efficient vascular systems, are vulnerable to extreme weather events, such as prolonged droughts, which can lead to the collapse of water and nutrient transport, affecting their survival (Araújo et al., 2024; Barros et al., 2019). This vulnerability is particularly worrying given that there is still a lack of large-scale studies on the impact of climate variations on the survival and occurrence for these large trees and species over time, especially *Dinizia excelsa*, the tallest in the Amazon. This gap limits our understanding of how climate factors may affect the stability and resilience of these critical forest components under future climate change scenarios.

Most studies focus on vegetation changes at regional scales and on individual analysis of species of various sizes (Klipel et al., 2022; Leão et al., 2021), often using data obtained from inventories or repositories such as the Global Biodiversity Information Facility - GBIF. Although these sources are valuable, they do not provide specific information about emergent trees, especially those exceeding 60 meters in height and growing in remote and difficult-to-access areas. In this context, the use of LiDAR (Light Detection and Ranging) data has proven essential for detecting and mapping forest vertical structure, enhancing our understanding of biodiversity and carbon stocks, and offering fundamental data for ecological and climate models (Asner et al., 2010; Coomes et al., 2017; Dubayah et al., 2020). By identifying height and distribution patterns of emergent trees, LiDAR enables us to understand how environmental factors influence the distribution of these giant trees, something difficult to achieve with traditional field methods due to the vastness and inaccessibility of the Amazon forest (Carvalho et al., 2023). Moreover, few studies specifically analyze how the tallest trees in the forest respond to climatic variations, such as changes in temperature and precipitation, especially in the context of global warming (Esquivel-Muelbert et al., 2019). With available forest inventory and LiDAR data, our research aims to address these issues, providing a more comprehensive analysis of how climate factors shape the survival, occurrence, spatial distribution, and habitat suitability of Amazonian giant trees, particularly under projected scenarios of extreme climate change.

Climate niche modeling emerges as a promising tool to explore these dynamics (Guisan et al., 2017; Scheffer et al., 2018; Zurell et al., 2024), allowing us to predict how the potential distribution of these trees may change under different climate variations and scenarios, as well as identify areas that may serve as future refuges for many giant species (De Lima et al., 2023). Given the climatic information on the Amazon critical inflection point (Flores et al., 2024), we can infer that minimum water availability and thermal stability may play determining roles in maintaining giant tree populations and, above all, that changes in these variables may result in a contraction of their distribution areas in the coming decades in the Amazon biome. Furthermore, we conjecture that, under more severe climate change scenarios, there may be a reduction in potentially suitable areas throughout the Amazon basin, possibly resulting in in habitat fragmentation for these trees (Flores et al., 2024; Nobre et al., 2016).

Here, we seek to provide a detailed overview of the vulnerability of giant Amazonian trees to climate change, integrating ecological niche models and remote sensing tools to identify the main factors underlying their distribution over time and space. More specifically, we intend to analyze the effects of the leading climate drivers on the occurrence and ecophysiological processes of the tallest trees obtained by LiDAR and of two large species inventoried in the Amazon, *Goupia glabra*, the most abundant large species (dbh ⩾ 70 cm) and *Dinizia excelsa* - the second most abundant large species and the tallest ever recorded in the Amazon (de Lima et al., 2022; Gorgens et al., 2019; Lewis et al., 2017). The expected results can contribute to the development of public policies and management strategies that ensure the conservation of these trees, which are essential for the climatic and ecological stability of the Amazon biome and provide a scientific framework for global initiatives to mitigate the effects of climate change.

## 2 Material and Methods

### 2.1 LiDAR data collection and occurrences of larger species

For this study, data on the occurrence of the tallest trees were obtained from the processing of the point cloud generated by airborne LiDAR mapping in 900 transects randomly distributed throughout the Amazon biome (Figure 1). The survey was carried out between 2016 and 2018 and is part of the EBA (Biomass Estimation in the Amazon) project developed by the National Institute for Space Research (Gorgens et al., 2019; Ometto et al., 2023). The sensor was the LiDAR HARRIER 68i, attached to a CESSNA model 206 aircraft. The scanning angle was 45 degrees, and the flight altitude was approximately 600 meters. Four returns per square meter form the point cloud. Each transect is represented in strips of 12.5 × 0.3 km, with an average area of 375 hectares. From the LiDAR points classified as ground, a Digital Terrain Model (DTM) was generated, which represents the ground elevation without the influence of vegetation. The Digital Canopy Height Model (CHM) was obtained by subtracting the DTM from the Digital Surface Model (DSM), including the total vegetation height. The CHM, therefore, represents the height of vegetation above the terrain. See more details at Zenodo Repository.

**Figure 1:**
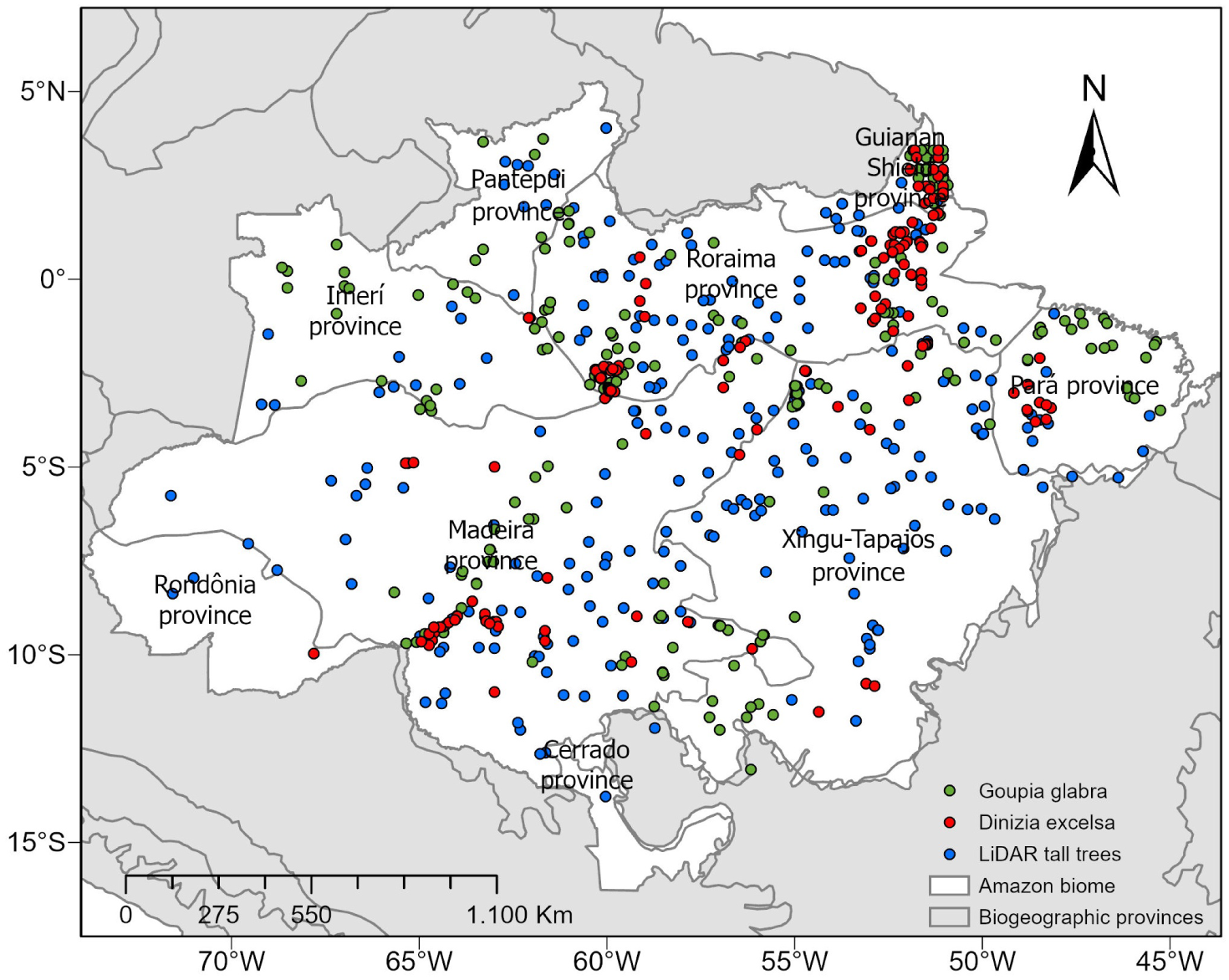
Map of the study area in Amazonia showing the distribution of the tallest trees and the location of the LiDAR samples. The green, red and blue dots represent the occurrences of *Goupia glabra*, *Dinizia excelsa*, and tall trees, respectively. The map also highlights the central biogeografic provinces of the Amazon such as the regions of Roraima, Guiana Lowlands, Pantepui, Imeri, Madeira, Xingu-Tapajós, and Rondônia, as well as the transition with the Cerrado biome to the south. The spatial context includes political boundaries and hydrography to better visualize the distribution of the samples in the Amazon biome.

A segmentation approach based on a 100-meter moving window was used to identify tree height and canopy metrics in the landscape (Roussel et al., 2024; Silva et al., 2022). This window spanned the CHM, and local canopy height maxima were extracted at each location. The local maxima detected by the moving window were interpreted as the points indicating the crowns of the tallest trees. The local tree crowns were grouped into three vertical strata: heights *<* 40 m, heights ⩾ 40 m and *<* 60 m, and heights ⩾ 60 m. The coordinates and heights of these trees were stored for later analyses. To model the niche of taller trees, we considered a giant tree to be one with a height ⩾ 60 m and defined a unique string for each transect with the occurrence (presence and absence) of trees in this section. This process was conducted based on a thorough analysis of all 900 mapped transects, and it resulted in a sample of 313 transects with the presence of at least one tree with a height ⩾ 60 m (blue dots in Figure 1, approximately 35% of the total) and 587 transects with the actual absence of taller trees distributed throughout the biome.

We also compiled a database at the species level, with georeferenced occurrences for the regionally known tree species Angelim vermelho – *Dinizia excelsa* and Cupiúba – *Goupia glabra*, obtained from forest inventories (de Lima et al., 2022) and the Global Biodiversity Information Facility - GBIF (www.gbif.org), using the R package ‘rgbif’ 3.7.5 (Chamberlain & Boettiger, 2017). These species were chosen given their gigantism characteristics, ecological importance, broad geographic distribution, and abundance of individuals with dbh ⩾ 70 cm in the Amazon (de Lima et al., 2022; Lewis et al., 2017).

Data with uncertain coordinates, missing latitude or longitude, marked as NA (not available), were eliminated using the ‘dplyr’ 1.0.10 (Wickham et al., 2023) and ‘tidyr’ 1.3.0 (Wickham et al., 2024) packages. To minimize spatial autocorrelation, duplicate occurrences within a 5 km radius were removed using the R package ‘spThin’ (Aiello-Lammens et al., 2019). This refinement yielded 213 georeferenced occurrence records for *Dinizia excelsa* and 568 for *Goupia glabra* across the Amazon basin. For each species, we defined an accessible area by applying a 100 km buffer around each occurrence point. Subsequently, we used the locations of field plots where the species did not occur, and 1000 background points were randomly generated within ten 10 × 10 km grid cells (Zurell et al., 2020), to mitigate potential biases from missing data during model calibration. There are different approaches to creating background or pseudo-absence data (Barbet-Massin, 2012; Kramer-Schadt et al., 2013; Phillips et al., 2009), although there is still room for further development in this field, and more precise recommendations for users would certainly be helpful (Zurell et al., 2020). Finally, we filtered the occurrence data by selecting the presence points for the species and for the set of tall trees (at least one larger tree mapped by LiDAR, inventory or gbif in each plot) and absence (no larger tree) within each transect or plots and assigning a binary string (1 and 0) using the ‘spThin’ package.

All mapped transects and points of occurrence of the species cover eight different ecoregions, as proposed by Morrone (2014). This biogeographic definition seeks to divide the Neotropical territory into areas with a common biogeographic history characterized by a particular set of species and evolutionary processes. Furthermore, we sought to understand the concept of niche or the suitable climatic conditions for potential zones and/or areas of occurrence of the giant trees (Scheffer et al., 2018; Scherrer et al., 2021). The occurrence data were then linked to spatially explicit and variable remote sensing (GIS) data to explore the underlying mechanisms that potentially govern the occurrence patterns of the tallest trees in the Amazon rainforest. The wide distribution of our sampling points (in number and distribution of plots and LiDAR transects) provides a substantially representative sampling effort to the larger trees of vegetation. It ensures that any uncertainty in the location of the samplings or minor changes in the forest area of the Amazon region hardly changed mean values or estimates of the probability of occurrence of the larger trees in the biome.

### 2.2 Acquisition and preprocessing of climate variables

We developed the spatial models using 19 bioclimatic covariates from WorldClim (www.worldclim.org) with a resolution of 2.5 arc minutes (Fick & Hijmans, 2017). We employed the Variance Inflation Factor (VIF) to avoid issues related to multicollinearity and mitigate the effect of high spatial correlation. We then eliminated the least significant bioclimatic variable from each pair with an absolute correlation greater than 0.8 using the R package ‘usdm’ (Naimi, 2023). We took special care when dealing with quarterly variables, ensuring they were considered carefully to avoid any undue influence on the modeling (Booth, 2022). Furthermore, we excluded variables with a VIF greater than a threshold of 10 from subsequent analyses. As a result, nine variables contribute to the set of tallest trees obtained by LiDAR and to the occurrence datasets of *Dinizia excelsa* and *Goupia glabra*, being: (1) Bio 3 - Isothermality; (2) Bio 4 - Temperature seasonality (standard deviation X 100; (3) Bio 5 - Maximum temperature of warmest month; (4) Bio 8 - Mean temperature of wettest quarter; (5) Bio 12 - Annual precipitation ; (6) Bio 13- Precipitation of wettest month; (7) Bio 14-Precipitation of driest month; (8) Bio 18 - Precipitation of the warmest quarter; and Bio 19 - Precipitation of the coldest quarter (see supplementary information table 2).

All selected spatial covariates were preprocessed using QGIS (Version for Windows - 3.34 LTR) and R 4.2.1 software (R Core Team, 2023 for Windows) and were then reprojected to the coordinate system of each sampling point area for calibration of the spatial model and to optimize the accuracy of the final figures and map areas. The first preparatory step in creating the spatial prediction was to assemble a data matrix with each sampling point’s location (longitude and latitude). Then, all the localized points were plotted on the stacked raster files of the geospatial covariates one by one. The geospatial metrics were extracted at each sample point using the *raster::extract* function of the ‘raster’ package (Hijmans et al., 2021) in R. This information was stored and saved in a final dataframe and used as predictor variables in the model (Figure 2).

**Figure 2:**
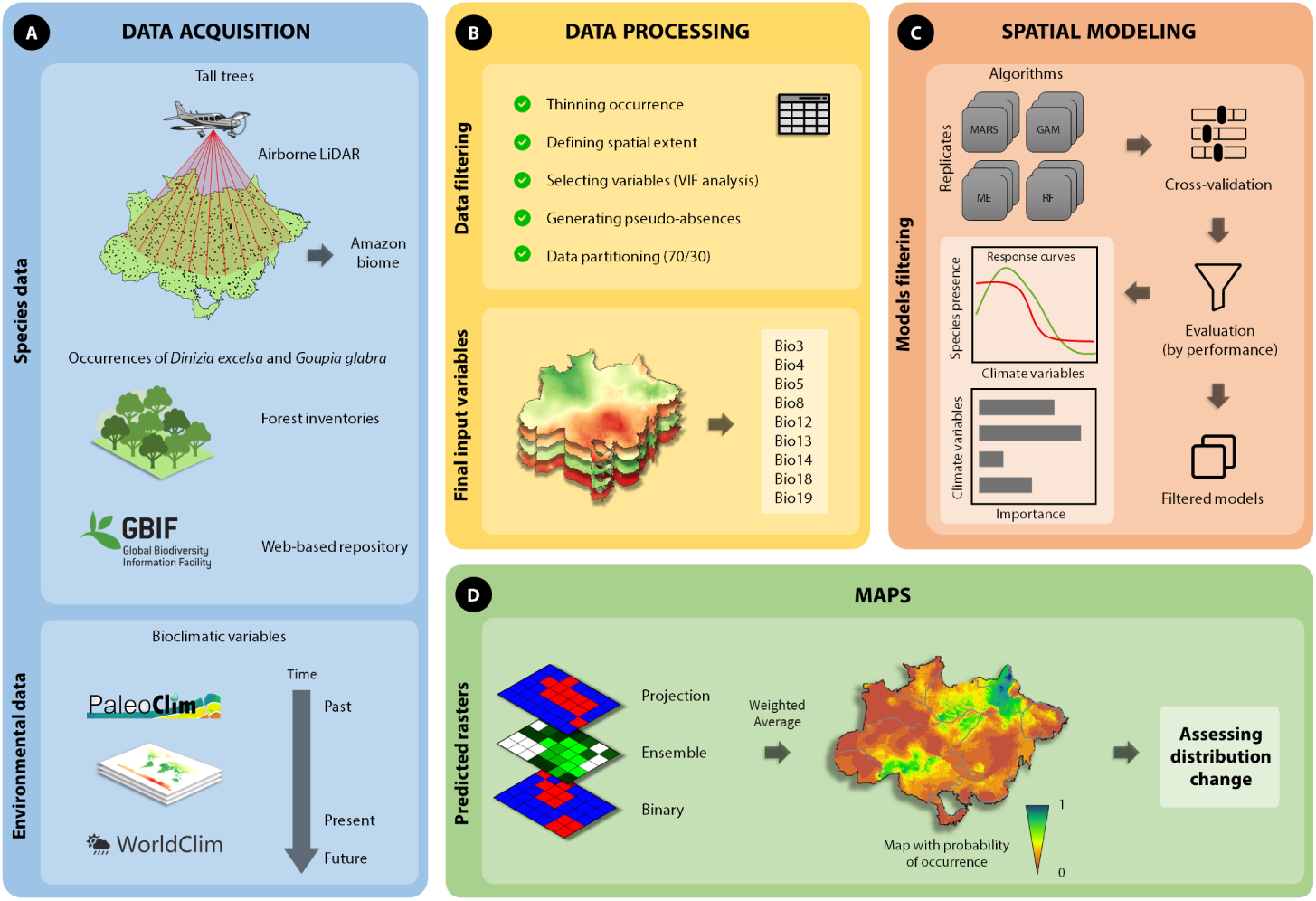
Methodological flowchart for mapping giant trees in the Amazon, highlighting the steps of data acquisition, processing, and spatial modeling. (A) Data collection: Airborne LiDAR surveys are used to identify and record giant trees (height ⩾ 60 m) and location data, ensuring accuracy in the Amazon. Additionally, tree occurrence data are obtained from forest inventories and online repositories such as GBIF. Environmental variables, including bioclimatic data, are extracted from sources like WorldClim and PaleoClim. (B) Data processing: Occurrence data are filtered to reduce spatial redundancy, define the study area extent, select relevant environmental variables through variance inflation factor (VIF) analysis, generate pseudo-absences, and partition data (70% for training and 30% for testing). (C) Spatial modeling: Different modeling algorithms, such as MARS, GAM, ME, and RF, are applied, with cross-validation and selection of the best-performing models based on evaluation metrics. Response curves are analyzed to understand the relationship between giant tree density and environmental variables. (D) Map generation: Model projections result in probability maps of giant tree occurrence, considering different approaches such as individual projections, ensemble models, and binary maps. The analysis of these maps allows for assessing changes in the spatial distribution of giant trees in the Amazon.

### 2.3 Climate niche and ensemble forecasts

We analyzed the potential occurrence of tall trees and the two species through climatic niche modeling. Climatic niche is a subset of the Ecological Niche Modeling concept, focused exclusively on climatic conditions (Guisan et al., 2017; Petitpierre et al., 2012; Soberón & Nakamura, 2009). Climatic niche models are widely used to predict the potential distribution of species, mainly in response to climate change. These models use climatic variables to estimate where current and future conditions may be suitable or unsuitable for the survival of a species (Guisan et al., 2017; Wiens et al., 2009). In this approach, we used the dataframe with the tall trees and species (*Dinizia excelsa* and *Goupia glabra*) occurrence (presence and absence) and the bioclimatic metrics selected in the VIF. For modeling with the occurrence of tall trees, we used the R package ‘sdm’ (Naimi & Araújo, 2016), selecting two algorithms from a total of 19 tested for this approach (Raes & Aguirre-Gutiérrez, 2018), namely MARS (Multivariate adaptive regression splines) and MaxEnt (Maximum Entropy).

For the species, we selected the three best algorithms: Generalized Additive Models (GAM), Maxent, and RandomForest (RF). MARS is used to model the nonlinear interactions between the predictor variables and the occurrence of giant trees. The algorithm builds a model based on splines that fit different linear functions to data intervals, efficiently capturing complex variations (Elith & Leathwick, 2009; Valavi et al., 2022). Maxent, one of the most widely used algorithms for species distribution modeling, was used to predict the probability of occurrence of giant trees by maximizing the entropy of the distribution of presences conditioned on climatic variables (Phillips et al., 2006). GAMs extend GLMs by allowing relationships between the predictor and response variables to be modeled nonlinearly using smooth spline functions. Despite their ability to model complex relationships, GAMs maintain interpretability, allowing them to visualize how each predictor variable affects the response quickly. The Random Forest (RF) algorithm (Cutler & Wiener, 2022) is a machine learning method that detects global trends present in data using a substantial ensemble of decision trees to predict tree occurrence in the uppermost strata of the Amazon rainforest canopy using the nine climate covariates. The RF algorithm applies the general bootstrap aggregation technique (bagging) with a modified tree learning algorithm that selects a random subset of the features at each candidate split in the learning process (Liang et al., 2022).

The data were randomly divided into a training data set (70%) and a test data set (30%). 100 replicates were performed for each model using resampling methods for subsampling. Sampling is done randomly and stratified to maintain the proportion of presences and absences in each subset. After fitting, the model is applied to the test set to predict the occurrences of presences and absences. With the predictions, we calculate the following metrics:

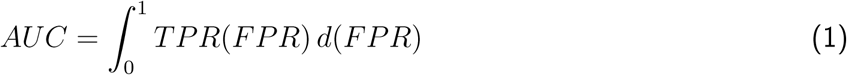

where:

- *TPR* is the True Positive Rate, also called sensitivity or recall;
- *FPR* is the False Positive Rate.

The True Skill Statistic (TSS) metric is calculated as:

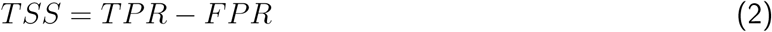

where:

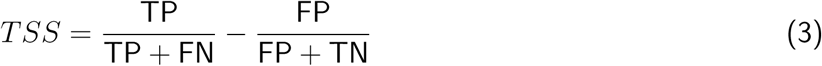

where:

- 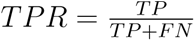 represents the True Positive Rate;
- 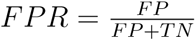 represents the False Positive Rate;
- *TP* is the number of true positives;
- *TN* is the number of true negatives;
- *FP* is the number of false positives;
- *FN* is the number of false negatives.

After all iterations, performance metrics are calculated for each replicate and then averaged to assess the overall model performance. Variability across validation runs is also analyzed, estimating model robustness and stability. Finally, we use the TSS-maximizing threshold (TSS ⩾ 0.70) to convert the probability of occurrence values into presence/absence predictions for binary transformation. The TSS-maximizing threshold approach is suitable because it produces the same threshold using presence-absence or presence-only data through sensitivity analysis (Guisan et al., 2017). Sensitivity analysis goes beyond evaluating each algorithm in isolation.

Next, we use the ensemble prediction procedure to obtain final models to reduce uncertainty between algorithms. This approach combines predictions from individual models to produce a final prediction. The ensemble models were predicted for past, current, and future climate conditions at a resolution of 2.5 arc-sec. Ensembles can reduce variability in predictions and minimize the risk of overfitting by leveraging the strengths and mitigating the weaknesses of different algorithms. In this study, we used Weighted Average; for example, instead of giving each model the same weight, we assigned different weights based on how well each model performed during validation. Models with better performance were given more weight in the final prediction. The weight was assigned using the maximized individual TSS values of each model.

Partial dependence plots were generated to visualize and interpret the relationship between each predictor variable and the probability of occurrence of these trees, keeping all other variables constant. This type of analysis allows us to examine the marginal effect of a specific variable, providing a clear view of how each environmental factor influences the distribution of tall trees, which is crucial for ecological niche models. In ecological niche modeling, partial dependence analysis is particularly valuable because it provides an understanding of the complex interactions between species and environmental gradients, facilitating the interpretation of models and helping to validate hypotheses about climate effects.

### 2.4 Assessing projections in climate change scenarios

We conducted these analyses, and we made predictions of the occurrence of the larger trees based on the climate conditions reconstructed for the past using the package ‘rpaleoclim’ (Roe, 2023) and projected for the future using package ‘geodata’ (Hijmans et al., 2024). For the past climate variables, we used the bioclimatic variables selected in the VIF reconstructed by the global general circulation model CCSM3 (Community Climate System Model ver. 3) for the Pleistocene: late-Holocene, Meghalayan (4.2-0.3 ka), which considers the chronological subdivision of the Holocene, more specifically the late Holocene, which began about 4,200 years ago (4.2 thousand years ago, or “ka”) and extends to approximately 300 years before present (0.3 ka) (Fordham et al., 2017). This division was established by the International Commission on Stratigraphy to describe a time interval marked by significant climate change in various parts of the world, such as a major global drought that affected several civilizations (Roe, 2023). This process was necessary because we assume that the larger trees in the biome may also be of advanced age. Therefore, if a giant tree or species can be long-lived, the climate that led to its regeneration, development, and distribution probably dates back much earlier (Guisan et al., 2017; Zurell et al., 2024). This is especially true if we assume that giant trees are centenarians and that the climate between 1720 and 1900 would have influenced their occurrence; therefore, it may coincide with the time of the aforementioned period.

For future climate conditions, we consider the scenarios projected for the epochs 2061–2080. These data were obtained from the CMIP6 general circulation model (Coupled Model Intercomparison Project - phase 6: BCC-CSM2-MR, CNRM-ESM2-1, and MIROC6; WorldClim v.2.1) of the Intergovernmental Panel on Climate Change (IPCC) Sixth Assessment Report (AR6) (Fick & Hijmans, 2017). This global climate model was selected based on its recent performance in case studies for modeling ecological niches, potential distribution, and habitat suitability for many plant species in tropical ecosystems (La Montagna et al., 2023; Leão et al., 2021). We additionally used shared socioeconomic pathways (SSPs) with low and high emission scenarios, namely SSP1-2.6 (optimistic) (van Vuuren, Stehfest, et al., 2011) and SSP5-8.5 (pessimistic) (Riahi et al., 2011), respectively. These SSPs were designed to describe various amounts of greenhouse gas emissions and potential future radiative forcings (van Vuuren, Edmonds, et al., 2011).

Finally, for each raster file generated by the ensemble predictions, we estimated absolute and percentage values of area gain and loss compared to the current distribution of tallest trees across the biome. In this step, we employed GIS tools for Map Algebra in QGIS to reclassify the raster files by probability of occurrence. We used spatial analysis tools to convert them to polygons. Soon after, in the reclassified file with defined polygons, we created a new column of attributes and calculated the sum of the areas in km2 of each class and the probability of occurrence and climatic niche of the tallest trees. To enhance the visualization of future projections, habitat suitability ratings were categorized into five distinct classes: very low suitability, low suitability, moderate suitability, good suitability, and high suitability. The analyses incorporated both absolute values in square kilometers (km^2^) and percentage changes relative to current conditions, allowing for a comprehensive assessment of the shifts in suitable habitats under future climate scenarios.

## 3 Results

### 3.1 Assessing Model Performance

The MAXENT and MARS algorithms suggest a feasible performance in habitat suitability projections for taller trees obtained by LiDAR. After validation with subsampling, the results revealed reliable performance metrics for each algorithm (overall average AUC = 0.84). Maxent demonstrated an average training AUC of 0.862, followed by MARS with 0.832. These values highlight the considerable discriminative ability of each model in separating classes of probability values for the occurrence of taller trees at the biome scale. The sensitivity analysis of the models used to predict the potential occurrence and suitable habitats for giant trees in the Amazon demonstrated an excellent performance of the algorithms. With a True Skill Statistic (TSS) value of 0.74 for the MARS model and 0.76 for the MAXENT model, both algorithms showed a high ability to distinguish areas of presence and absence of giant trees, with a slight superiority of MAXENT in terms of predictive accuracy.

For *Dinizia excelsa*, AUC values ranged from 0.82 to 0.85. The Random Forest model presented the highest AUC value (0.85), followed by GAM and MaxEnt, with 0.82. These values above 0.8 suggest that all methods have good predictive ability for this species, with Random Forest being the most accurate. The TSS was also highest in the Random Forest model (0.78), followed by MaxEnt (0.73) and GAM (0.72), reinforcing the superiority of Random Forest for this species. For *Goupia glabra*, AUC values are similar, ranging from 0.83 to 0.86. Random Forest again obtained the highest AUC value (0.86), indicating a moderate performance in predicting the distribution of this species, with GAM (0.75) and MaxEnt (0.73) presenting slightly lower values. In the case of TSS, the values were lower for *Goupia glabra*, with Random Forest presenting the highest value (0.71), followed by GAM (0.79) and MaxEnt (0.74).

### 3.2 Ideal climatic conditions for the occurrence of the larger trees

The analysis of the effects of climate variables on the occurrence of tall trees and the species *Dinizia excelsa* and *Goupia glabra* in the Amazon reveals important ecological patterns (Figure 3). Notably, areas influenced by Isothermality, maximum temperatures during the warmest month, and temperature seasonality are less likely to harbor larger trees. The negative effect of maximum temperature, a significant variable, becomes pronounced when it exceeds 32 °C, thereby limiting the suitability of the habitat for giant trees. Isothermality, which reflects daily temperature variation about annual temperature, shows an unimodal pattern with a negative effect on the occurrence of tall trees when values are above 75-80%. This effect suggests that moderate thermal stability contributes to the habitat suitability of these trees. This finding underscores the sensitivity of tall trees to thermal extremes, particularly in the Amazon forests, where such events can induce water stress and hinder growth.

**Figure 3:**
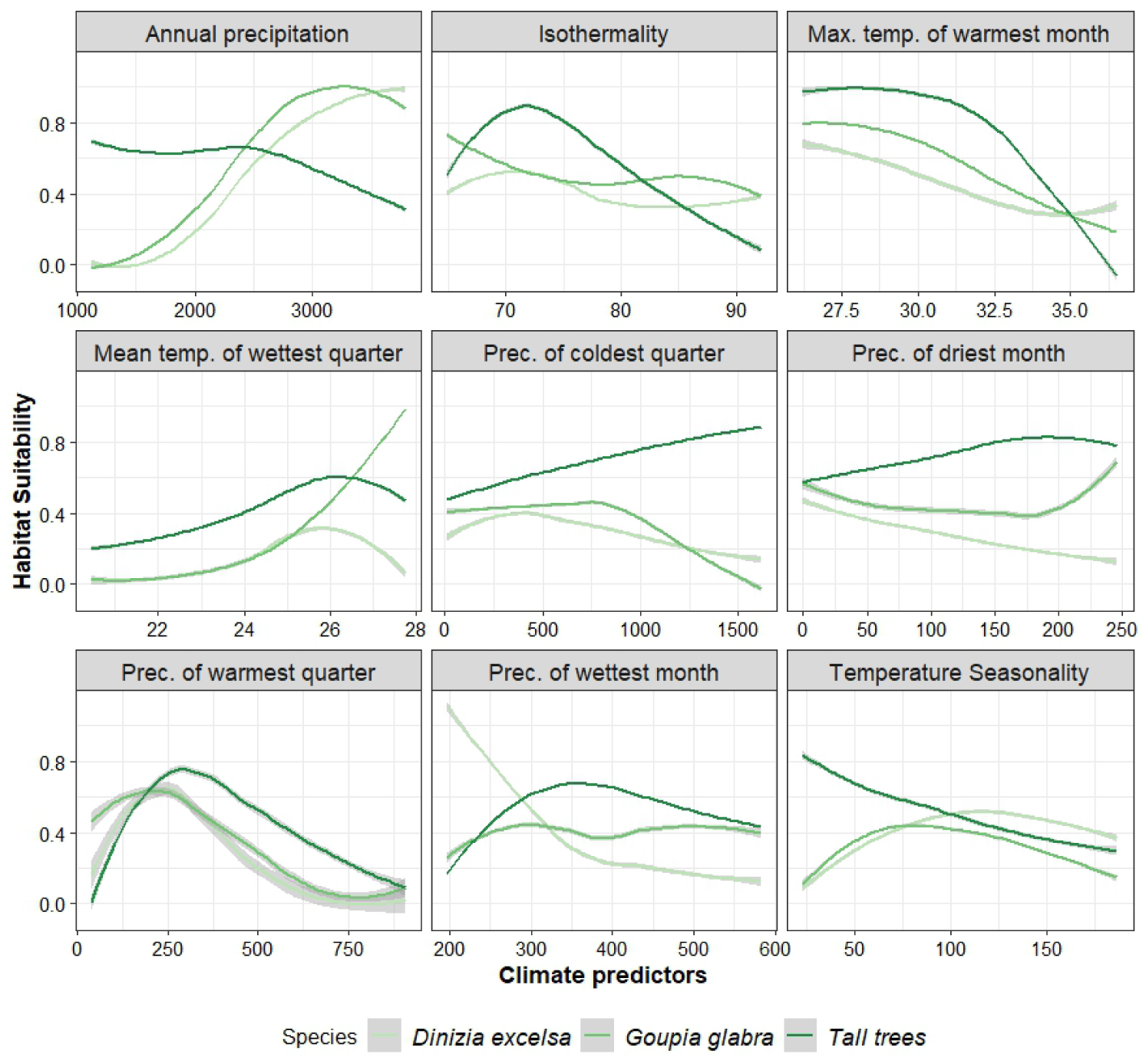
Partial dependence plots showing the relationship betwen habitat suitability and selected climatic variables for tall trees, *Dinizia excelsa*, and *Goupia glabra*. The plots were generated using an ensemble model to analyze how different climatic predictors influence habitat suitability for the studied species. The dark, medium, and light lines represent tall trees, *Dinizia excelsa* and *Goupia glabra*, respectively. These results highlight the ecological plasticity of the species in response to different climatic conditions and their ability to adapt to environmental variations.

On the other hand, areas with greater water availability throughout the year indicate a higher probability of giant trees occurring. Precipitation in the driest month, coldest quarter, and wettest month have a strong positive relationship with habitat suitability for tall trees, indicating that the continuous presence of water at different times of the year is essential. The average temperature of the wettest quarter, however, emerges as a key factor, indicating a preference for occurrence in areas where this variable is between 24-26 °C. This likely ensures a favorable environment for growth during the period of greatest precipitation, thereby underscoring the role of temperature in the Amazon’s biodiversity.

In addition, the analysis of the essential variables reinforces the critical environmental conditions for the occurrence of tall trees and *Dinizia excelsa* and *Goupia glabra* (Figure 4). Precipitation in the hottest quarter appears to be one of the most relevant for tall trees and *Dinizia excelsa*, indicating that places with around 500 mm of rain during this period offer ideal conditions. Isothermality is also essential (relative importance AUC = 0,25%), especially for tall trees and *Dinizia excelsa*, highlighting the importance of moderate thermal variation for the occurrence of these species. In contrast, *Goupia glabra* is more influenced by precipitation in the driest months and the coldest quarter, reflecting a greater tolerance to climatic variations than *Dinizia excelsa*. Finally, annual precipitation shows a multimodal effect, suggesting that intervals of 1500 mm and 2500 mm are favorable for the occurrence of these tall trees.

**Figure 4:**
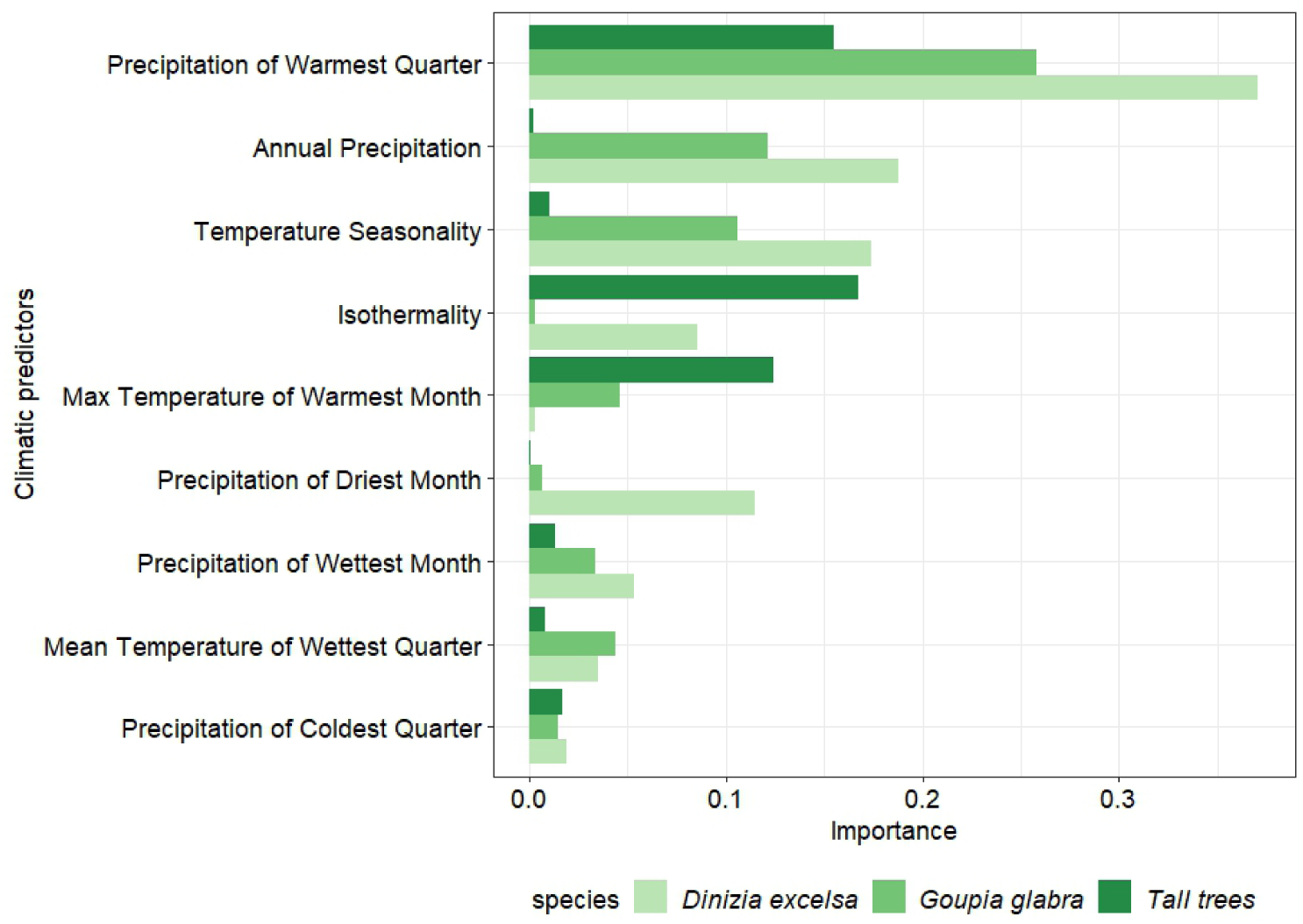
The most important selected climate variables in the ensemble model to predict the distribution of tall trees, *Dinizia excelsa*, and *Goupia glabra*. The bar graphs indicate the relative importance of each climate variable, highlighting the precipitation of the driest quarter, seasonal temperature, and isothermality, which are determinants for the presence of these species in specific regions of the Amazon. Each color represents a class of trees: tall trees (dark green), *Goupia glabra* (medium green), and *Dinizia excelsa* (light green).

### 3.3 Past, Current, and Future Prediction Maps

The analysis suggests that historical climatic conditions may have significantly influenced the occurrence of these trees in specific regions (Figure 5). Areas with the highest probability of occurrence, indicated by colors closest to 1 (green-blue), are concentrated in the northern and northwestern Amazon, especially in the Guiana Shield and Roraima provinces, which span approximately coordinates 3°S to 5°N and 60° to 50°W. These regions, characterized by a combination of stable isothermality and well-distributed precipitation throughout the year, appear to have offered ideal climatic conditions in the past for the establishment and growth of giant trees and the two species studied. In the Xingu-Tapajós province (5°S to 0°, 60° to 50°W), habitats were also largely suitable, although with slightly lower suitability than in areas to the north. In the Madeira province, located between 10° to 5°S and 65° to 55°W, past climatic conditions present an intermediate probability of supporting these species. On the other hand, in the southernmost areas of the Amazon, such as the province of Rondônia (approximately 15° to 10°S and 65° to 60°W), suitability decreases considerably. These patterns suggest that climatic variables, such as temperature seasonality and precipitation in the driest and wettest months, played a crucial role in shaping the climatic niches of these species and tallest trees in specific regions of the Amazon.

**Figure 5:**
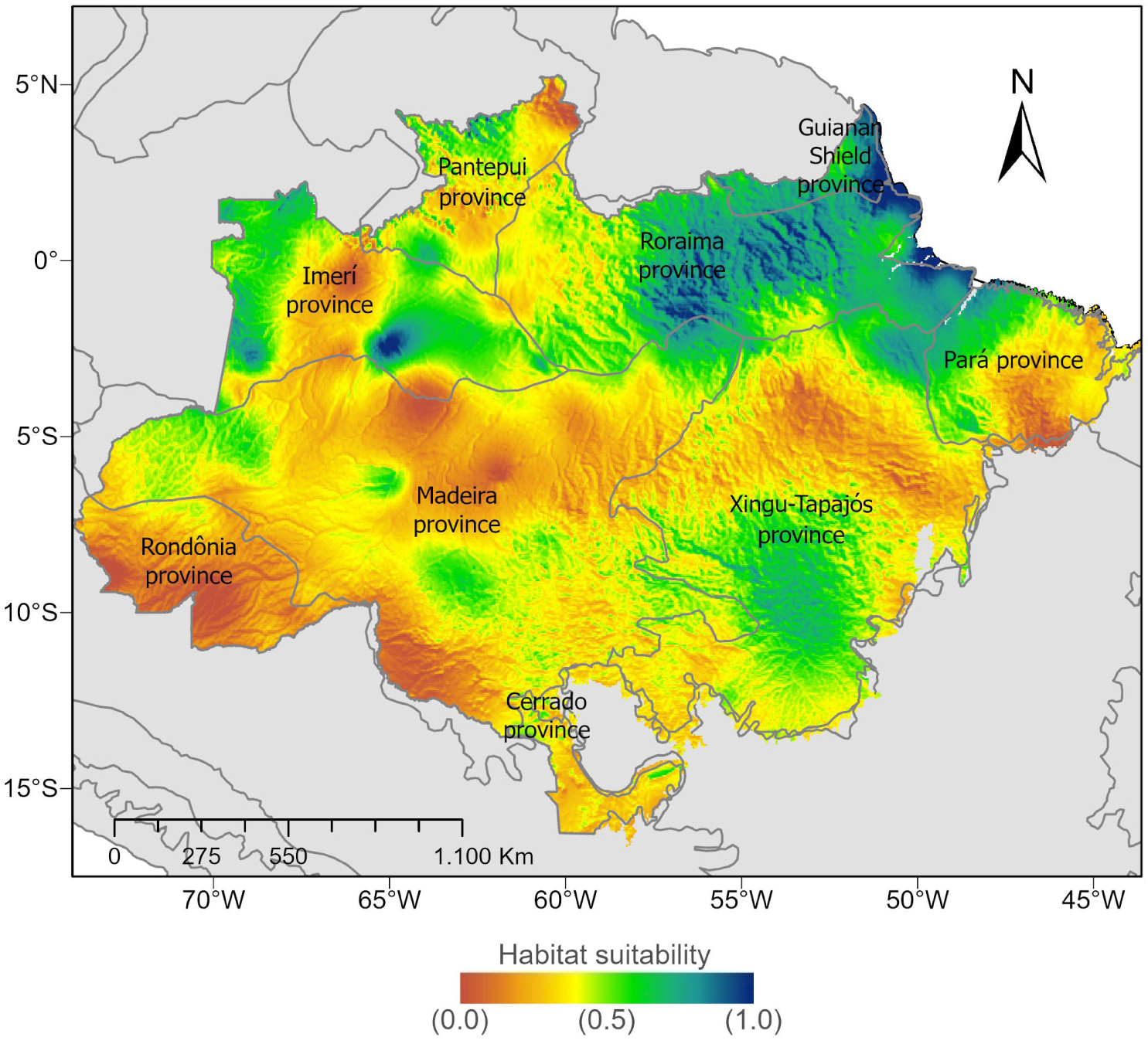
Map of past suitable habitats for giant trees in the Amazon using historical climate data and selected bioclimatic variables. This map reconstructs the distribution of past suitable habitats for giant trees, highlighting areas with climatic conditions that historically favored the growth of these trees. Areas in darker tones indicate a higher probability of occurrence.

Under current climatic conditions, suitable areas appear to be more concentrated in the northern and central parts of the Amazon, such as the Guiana Shields Province, Roraima Province, and the western portion of the Xingu-Tapajós Province (Figure 6). The Pará Province, which in the past had some areas of suitability, shows an increase in the concentration of green areas, indicating a shift in favorable climatic conditions in this region. This shift may result from variations in the Isothermality and average temperature of the wettest quarter, which are now more favorable for these trees. In general, despite the visible dynamism between provinces at different times, the general pattern of potential distribution and suitable habitats in the Amazon region remains stable in the Province of Roraima, with slight changes in the eastern portion and significant gains in suitable areas in the Provinces of Madeira and Xingu-Tapajós. However, in regions such as the Imeri and Pantepui Provinces, it appears to have decreased, suggesting changes in climatic conditions that have negatively affected the presence of these trees in these areas over time.

**Figure 6:**
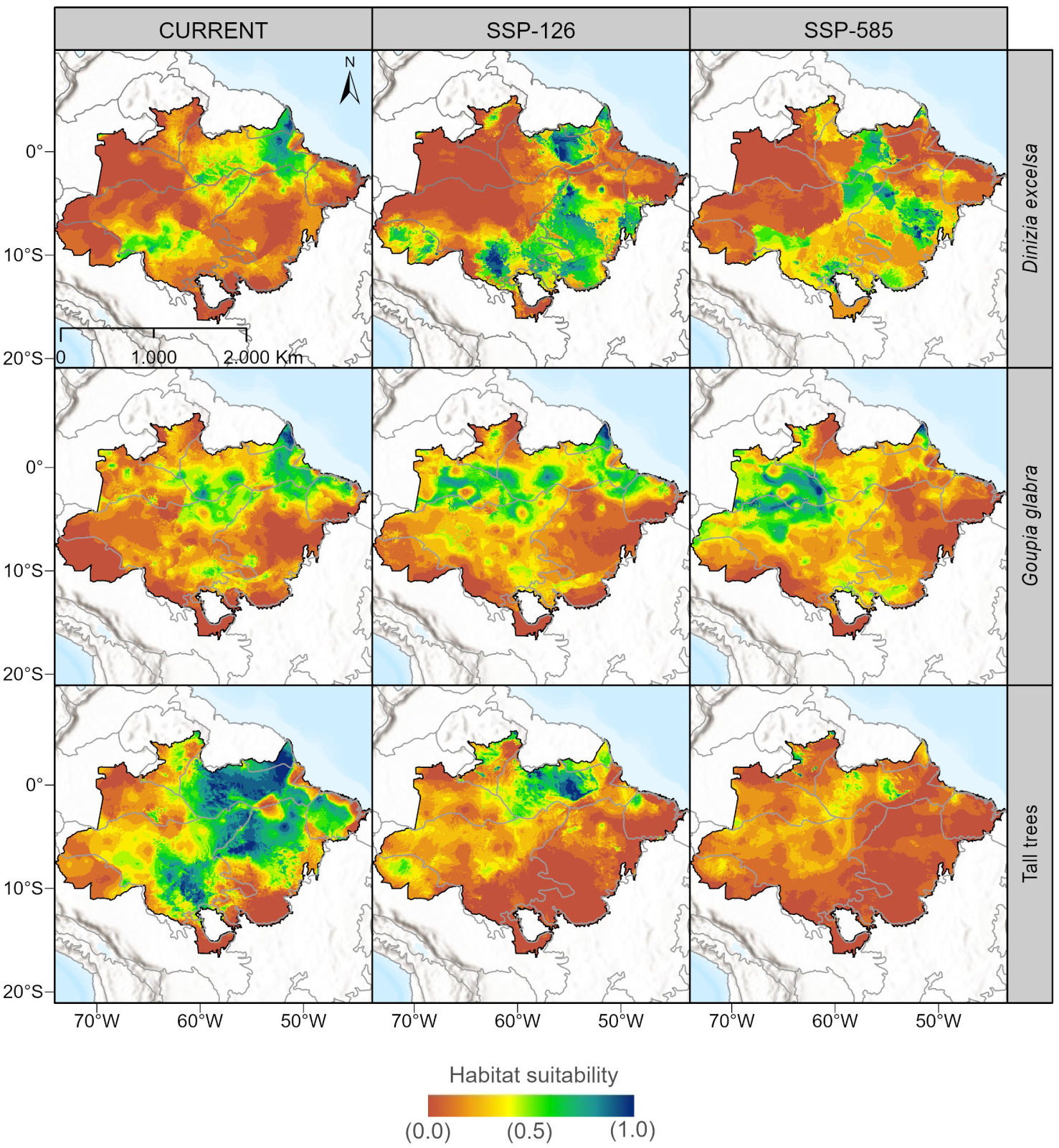
Potential distributions for *Dinizia excelsa*, *Goupia glabra* and tall trees under different future climate change scenarios (SSP1-2.6 and SSP5-8.5). The models project the potential distributions of these species considering climate emissions scenarios based on the latest IPCC report. The maps indicate how changes in temperature metrics, precipitation, and other predictors may influence the expansion or contraction of suitable habitats for these species and tall trees.

Overall, the ensemble model’s results indicate a substantial reduction in the spatial extent of suitable habitats for giant trees across the Amazon. The loss of potential distribution is expected to accelerate significantly by the late 2080s. The predicted distribution patterns under the different Shared Socioeconomic Pathway (SSP) scenarios exhibit similarities, particularly in the optimistic SSP1-2.6 scenario. However, under the pessimistic SSP5-8.5 scenario, the persistence of suitable environmental conditions for these species will be restricted to small, fragmented areas concentrated in specific biogeographic provinces.

*Dinizia excelsa* is predominantly distributed in the northern and central Amazon, with its highest concentrations in the Guiana Shield, Roraima, and the western portion of the Xingu-Tapajós Province. Future projections under SSP5-8.5 suggest a drastic contraction of suitable habitats, with high-suitability areas expected to decrease by 71%, corresponding to a loss of 82,286 km^2^, leaving only 145,180 km^2^ available by the 2061-2080 period. The provinces of Imeri and Pantepui are anticipated to be the most affected, where rising temperatures and decreasing water availability will impose significant environmental stress, leading to a severe decline in habitat suitability. By 2080, very low suitability areas are projected to expand by 89.75%, covering 1,717,325 km^2^, while low suitability areas will remain relatively stable at 1,142,503 km^2^. Moderate suitability habitats will experience a minor fluctuation, with a 5.85% gain. However, good suitability areas will shrink by 124,244 km^2^, representing a 32% decline. The most critical impact will be observed in high suitability areas, which will diminish to just 145,180 km^2^, reflecting a 71% reduction compared to current conditions. Under the SSP1-2.6 scenario, however, the Guiana Shield and Pará provinces are expected to experience a 15% increase ( 200,000 km^2^) in suitable habitats, offering potential climate refugia. Conversely, the southern Amazon, particularly in the arc of deforestation, will become increasingly unsuitable due to intensifying droughts and extreme temperatures, further threatening the persistence of Dinizia excelsa in these regions.

The current distribution of *Goupia glabra* spans much of the central and northern Amazon, with significant populations in the Guiana Shield and the Guiana Shield Province. Under SSP1-2.6, the species is projected to experience a 10% expansion in areas of intermediate suitability, particularly in the northern Amazon and the Guiana Shield, where climate conditions are expected to remain relatively stable. However, under SSP5-8.5, a substantial reduction of 30% ( 400,000 km^2^) in suitable habitat is projected, primarily affecting the Xingu-Tapajós and Roraima provinces, where increasing extreme temperatures and declining precipitation will create unfavorable conditions for the species. By 2080, very low suitability areas are expected to expand by 76.72%, covering 968,487 km^2^, while low suitability habitats will remain relatively unchanged at 1,296,042 km^2^. Moderate suitability areas will initially increase but are predicted to decline by approximately 262,744 km^2^. Good suitability habitats will undergo a moderate reduction of 85,529 km^2^. The most pronounced change will be in high suitability areas, which are projected to expand significantly under SSP5-8.5, reaching 251,639 km^2^, representing a 291% increase. This trend may be associated with the migration of suitable conditions toward more stable northern regions. The southern Amazon, particularly the Rondônia Province, is expected to become inhospitable for *Goupia glabra*, contributing to a fragmented habitat mosaic. This decline in suitable areas aligns with the projected savannization of the Amazon, where structurally complex forests are replaced by lower-diversity and less productive ecosystems, further exacerbating biodiversity loss and reducing carbon sequestration capacity.

Tall Trees exhibit a similar trend, with extreme reductions in suitable habitats across the Amazon. Under SSP5-8.5, the spatial distribution of highly suitable areas will be nearly eliminated, leaving only small, fragmented patches of viable habitat. The expansion of very low suitability areas is particularly striking, with a 274% increase, reaching 2,286,498 km^2^ by 2080. Similarly, low-suitability habitats will expand by 139%, affecting 1,432,150 km^2^. In contrast, moderate suitability areas will decline by 51%, restricting viable habitat availability. Good suitability areas will experience an even more dramatic loss, shrinking by 94%—equivalent to 688,410 km^2^—nearly eradicating these habitats. High suitability areas will be the most severely impacted, undergoing a 99% decline, losing 725,311 km^2^, leaving only 1% of the current distribution intact. The provinces of Roraima, the Guiana Shield, and western Pantepui will become the last strongholds for emergent trees, while the Madeira Province will see a sharp decline in occurrence probability. Meanwhile, a broad southern strip of the biome, aligned with the arc of deforestation, is expected to become almost entirely unsuitable for giant trees, reinforcing the trend of habitat degradation. (Figure 6 Table 1).

**Table 1:**
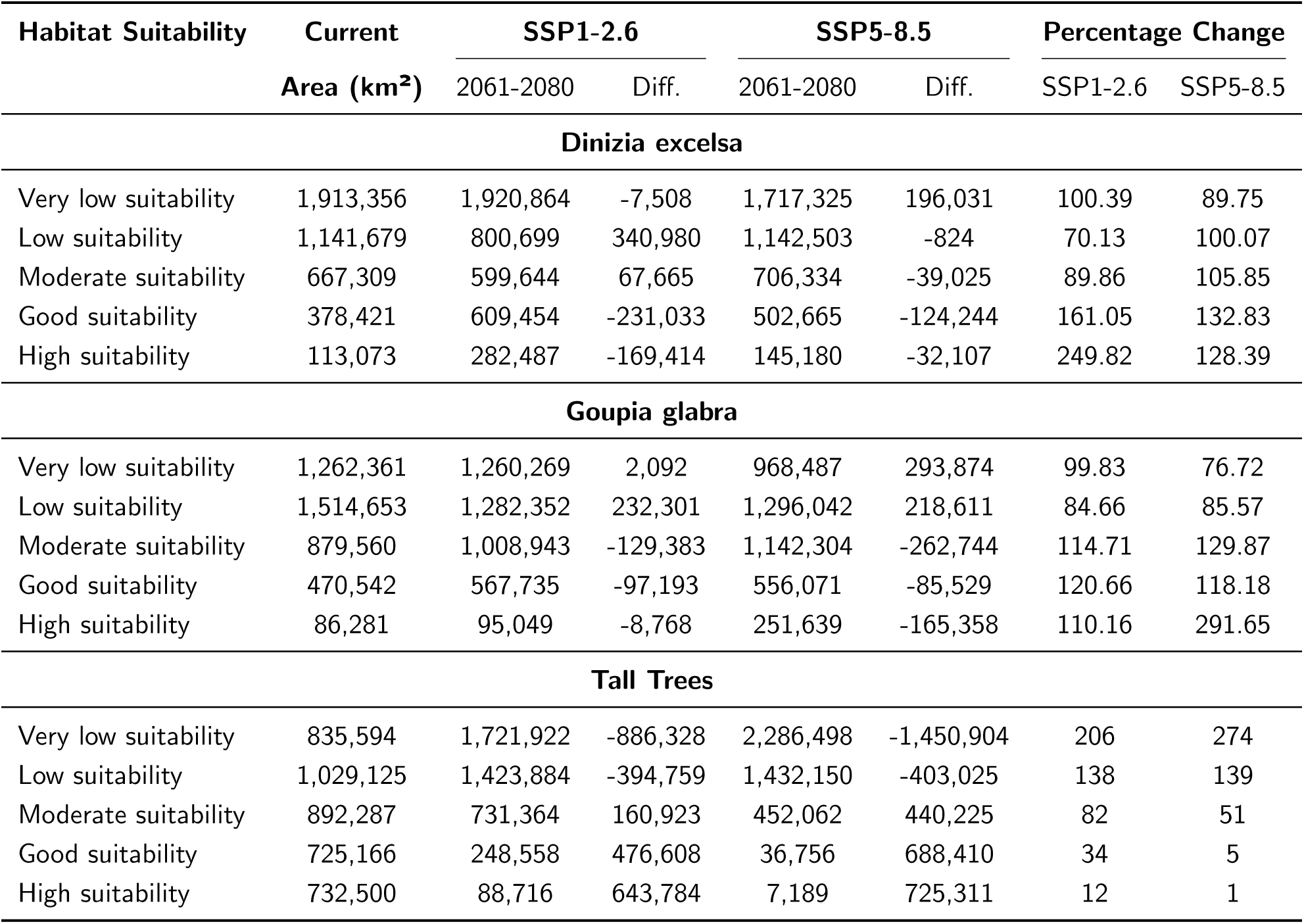
Projected changes in habitat suitability (area in km^2^) for *Dinizia excelsa*, *Goupia glabra*, and Tall Trees under climate scenarios SSP1-2.6 and SSP5-8.5 for the period 2061-2080. Values represent the total area in each category, differences from the current scenario, and percentage changes.

| Habitat Suitability | Current | SSP1-2.6 |  | SSP5-8.5 |  | Percentage Change |  |
| --- | --- | --- | --- | --- | --- | --- | --- |
|  | Area (km<sup>2</sup>) | 2061-2080 | Diff. | 2061-2080 | Diff. | SSP1-2.6 | SSP5-8.5 |
| **Dinizia excelsa** |  |  |  |  |  |  |  |
| Very low suitability | 1,913,356 | 1,920,864 | -7,508 | 1,717,325 | 196,031 | 100.39 | 89.75 |
| Low suitability | 1,141,679 | 800,699 | 340,980 | 1,142,503 | -824 | 70.13 | 100.07 |
| Moderate suitability | 667,309 | 599,644 | 67,665 | 706,334 | -39,025 | 89.86 | 105.85 |
| Good suitability | 378,421 | 609,454 | -231,033 | 502,665 | -124,244 | 161.05 | 132.83 |
| High suitability | 113,073 | 282,487 | -169,414 | 145,180 | -32,107 | 249.82 | 128.39 |
| **Goupia glabra** |  |  |  |  |  |  |  |
| Very low suitability | 1,262,361 | 1,260,269 | 2,092 | 968,487 | 293,874 | 99.83 | 76.72 |
| Low suitability | 1,514,653 | 1,282,352 | 232,301 | 1,296,042 | 218,611 | 84.66 | 85.57 |
| Moderate suitability | 879,560 | 1,008,943 | -129,383 | 1,142,304 | -262,744 | 114.71 | 129.87 |
| Good suitability | 470,542 | 567,735 | -97,193 | 556,071 | -85,529 | 120.66 | 118.18 |
| High suitability | 86,281 | 95,049 | -8,768 | 251,639 | -165,358 | 110.16 | 291.65 |
| **Tall Trees** |  |  |  |  |  |  |  |
| Very low suitability | 835,594 | 1,721,922 | -886,328 | 2,286,498 | -1,450,904 | 206 | 274 |
| Low suitability | 1,029,125 | 1,423,884 | -394,759 | 1,432,150 | -403,025 | 138 | 139 |
| Moderate suitability | 892,287 | 731,364 | 160,923 | 452,062 | 440,225 | 82 | 51 |
| Good suitability | 725,166 | 248,558 | 476,608 | 36,756 | 688,410 | 34 | 5 |
| High suitability | 732,500 | 88,716 | 643,784 | 7,189 | 725,311 | 12 | 1 |

## 4 Discussion

### 4.1 Underlying climatic drivers for the occurrence of larger trees

In tropical ecosystems, the climatic factors are fundamental drivers for the development and survival of tree species, especially those that dominate the highest vertical stratum of the forest. They act directly on the ecophysiological processes essential for productivity and maintenance in different environments. The Amazon biome, characterized by its vast diversity of habitats, is subject to climatic variations that can significantly affect the performance of the larger trees with high water and energy resource requirements. Therefore, analyzing the effect of climatic factors is crucial to understanding the distribution and occurrence of many giant species and, above all, to provide new insights to discuss the complex interactions between climate and ecosystem in the region.

Our analyses showed that Isothermality and the maximum temperature of the hottest month had adverse effects on the occurrence of the tallest trees and are between the most important variables associated with temperature in the spatial model (Figure 4). These variables can directly affect metabolism, transpiration rate, and photosynthesis. In simple terms, Isothermality expresses how constant or variable the temperature is over a daily or seasonal cycle relative to the annual variation. Figure 3 demonstrates that the suitability response to isothermality peaks around 70-75%, decreasing as thermal variations increase. More tall trees have a canopy exposed to different solar radiation levels throughout the day, which may increase the need to maintain a thermal balance (Kăspar et al., 2024; Manzi et al., 2024). Therefore, places with moderate Isothermality (close to 70%) are considered favorable environments for the occurrence of giant trees, as they contribute to maintaining a stable climate that can favor continuous growth in height and biomass (Gorgens et al., 2021). However, in areas with high Isothermality (less temperature variation throughout the year and day), the occurrence of taller trees may be affected, possibly because some species depend on seasonal temperature variations to regulate ecophysiological processes, such as changes in transpiration rate or resource allocation for defense against heat (Q. Song & Zhu, 2024). When Isothermality is high, these variation cycles are reduced, which can negatively impact their development in several ways, such as reduced photosynthetic efficiency, limitations in water transport, and reduced growth stimulation (Natale et al., 2023; Sexton et al., 2021; Wu et al., 2017).

Extreme heat stress, particularly the maximum temperature of the hottest month, can significantly limit tree growth by impairing photosynthetic efficiency and increasing transpiration rates (Anderson et al., 2021; Koch et al., 2004; Y. Song et al., 2014; Teskey et al., 2015). Prolonged exposure to these high temperatures can lead to the denaturation of critical enzymes such as rubisco, which are essential for photosynthesis (Natale et al., 2023; Wijewardene et al., 2021). Consequently, regions within the biome that experience significant temperature fluctuations and high isothermality levels may face increased water stress in tall trees, thereby reducing their growth potential. Under these conditions, trees often increase transpiration to cool down, which can also lead to substantial water loss, further exacerbating stress (Meinzer et al., 2001; Sadok & Lopez, 2020; Stephenson & Das, 2020). In areas where maximum temperatures exceed 34°C, models predict a marked decline in suitable habitats, as higher temperatures significantly lower the probability of tree occurrence (Figure 3). However, some trees exhibit adaptive mechanisms, such as reducing leaf area to mitigate heat stress (Miller et al., 2021; Sexton et al., 2021). This temperature variability may favor species adapted to extreme seasonal conditions, enhancing their functional plasticity in ecophysiological responses (Liu et al., 2019; Vinod et al., 2023). Tall trees, for instance, may adjust their canopy structure to minimize direct sun exposure and increase stomatal density to regulate water loss through evapotranspiration (Miller et al., 2021; Stevens et al., 2021; Still et al., 2022).

These results corroborate the curve of Temperature Seasonality, both for the set of tall trees mapped by LiDAR and for *Dinizia excelsa* and *Goupia glabra*. That is, areas with high seasonality are characterized by more significant thermal fluctuations, which can increase thermal stress in trees whose canopies have greater exposure to the sun (Manzi et al., 2024; Scheffer et al., 2018), especially in the hottest periods of the year. Temperature seasonality refers to the amplitude of thermal variation between different year periods, such as hot and cold seasons. This variability can directly affect larger trees’ nutrient absorption capacity and imposes significant physiological limitations, significantly when it exceeds the thermal tolerance threshold of the trees (Wahid et al., 2007). As a result, trees in regions with high thermal seasonality and high temperatures need to develop mechanisms that allow them to withstand significant temperature variations without suffering irreversible damage, such as adjustments in metabolism or structural changes in wood (Barros et al., 2019; Bhatla & Lal, 2023a, 2023b; Nievola et al., 2017). For taller trees, which already have a greater demand for energy and water due to their size (Liu et al., 2019), this elasticity in physiological adjustments includes the ability to maintain efficient water transport through the xylem even under high temperatures (Domec et al., 2008; Meinzer et al., 2001; Olson et al., 2018). However, the risk of embolism can increase dramatically due to the greater height of the water column that they need to maintain (Araújo et al., 2024; Fernández-de-Unã et al., 2023; Garcia et al., 2023; Oliveira et al., 2019). Despite these adverse effects, larger trees can adjust their growth cycle by allocating resources more efficiently across seasons to optimize light capture and minimize water loss during hotter and drier periods, and they can modify their architecture and root length and distribution (Banin et al., 2012; Iida et al., 2011; Meinzer et al., 2001; Sexton et al., 2021).

On the other hand, larger trees are susceptible to water availability, mainly due to their high water requirements (Maréchaux et al., 2015). For example, in areas where the average rainfall in the warmest quarter is 300 mm, the minimum water availability required by larger trees is crucial to maximize nutrient uptake and maintain their physiological processes at high levels (Bartholomew et al., 2022; Y. Song et al., 2014; Zweifel et al., 2006). In the spatial model, this variable was the most important for the species *Dinizia excelsa* and *Goupia glabra* and was the second most important for the set of tall trees (Figure 4). These results are consistent with the response curve of the average temperature of the wettest quarter (a period that coincides with the time of most excellent water availability and least water stress) varying around 24°C to 26°C (as shown in Figure 3). During the period of these variables in the year, trees can optimize water use without losing metabolic efficiency due to extreme temperatures (Gwitira et al., 2014; Teskey et al., 2015; Vinod et al., 2023). The temperature of the wet quarter can directly influence sap flow within the vascular system and can also promote favorable conditions in tree phenology, such as the production of new leaves, flowering, and fruiting (Brearley et al., 2007; Pereira et al., 2024; Schaik et al., 2003; Schöngart et al., 2002; Zurell et al., 2024), which facilitates dispersion and distribution over time and space for taller trees and large species (Slik et al., 2024). Moderate temperatures during the wet quarter can contribute to the fact that water transport from the roots to the leaves is efficient, avoiding the risk of xylem embolism, which is more likely under conditions of heat stress or water deficit (Araújo et al., 2024; Bittencourt et al., 2020; Meinzer et al., 2001; Olson et al., 2018).

A combined effect can also be seen when Isothermality is moderate. For example, trees face less daily heat stress, which facilitates the regulation of transpiration and water balance (Manzi et al., 2024; Sadok & Lopez, 2020). During the wet quarter, moderate temperatures and high water availability allow trees to maximize growth without compromising their ability to adjust to extreme climate variations (Kăspar et al., 2024; Reich et al., 2015). Giant trees, which have a complex vascular system and a high water requirement, benefit from these stable conditions (Esquivel-Muelbert et al., 2019). With moderate Isothermality, trees experience a minor temperature variation between day and night, which helps regulate photosynthesis and nocturnal respiration (Becklin et al., 2021; Larjavaara, 2014; Natale et al., 2023). During the wettest quarter, this thermal stability, combined with the moderate average temperature, allows trees to maintain a high rate of photosynthesis without wasting resources on excessive respiration (Mathur et al., 2014; Miller et al., 2021). However, in periods with water deficit conditions and high temperatures, thermal and water stress combine, forcing trees to close their stomata to avoid excessive water loss, which decreases photosynthesis and negatively affects biomass production (Anderson et al., 2021; Sadok & Lopez, 2020). Precipitation during the driest month is a critical indicator of water availability during the most significant water deficit periods. Due to their large leaf area and need to support large-volume tissues, giant trees are highly dependent on soil water availability (Bartholomew et al., 2022; Spanner et al., 2022; Terra et al., 2018). During the driest month, the lack of precipitation can significantly reduce transpiration and directly impact water transport to the highest branches of the canopy (Liu et al., 2019; Sadok & Lopez, 2020). Although these trees can develop deep roots to access water reserves in deeper soil layers (Esquivel-Muelbert et al., 2019), prolonged water stress can cause early leaf senescence, xylem embolism, and, in more severe cases, mortality (Aleixo et al., 2019; Allen et al., 2010; Skiadaresis et al., 2021).

In general, precipitation regimes (wettest month precipitation and coldest quarter precipitation) determine the health and distribution of giant trees in the Amazon. Adequate precipitation (around 1500-2500 mm annually) provides constant soil moisture to sustain the biomass of these trees. In Figure 3, habitat suitability increases with precipitation in the wet months, significantly above 400 mm, indicating that the distribution of giant trees is strongly linked to water availability during these periods. However, although the Amazon is known for its high levels of precipitation, excessive precipitation above 2800 mm per year can create unfavorable conditions for the occurrence of tall trees (Figure 3). Large trees, especially in the Amazon, depend on a series of balanced ecophysiological factors to maintain their growth and vigor. When precipitation exceeds a specific limit, adverse effects can arise, directly affecting the health and growth of these trees. For example, excessive precipitation can lead to soil saturation, reducing the oxygen available to tree roots (Furtak & Wolińska, 2023; Sprunger et al., 2023). High precipitation can accelerate the leaching process, where essential nutrients for tree growth, such as calcium, potassium, and magnesium, are carried to deeper soil layers or watercourses. This can result in nutrient-poor soils in the surface layers, where tree roots absorb most of their resources (Giehl & Wirén, 2014; Rashmi et al., 2017). Environments with high humidity and constant rainfall can also favor the proliferation of fungal diseases, especially on tree foliage and roots (Velásquez et al., 2018). Fungi, such as those that cause root rot, thrive in moist conditions, compromising the integrity of the root system and increasing the tree’s susceptibility to falling or severe infections (Birch et al., 2023).

### 4.2 Impact of climate change on the occurrence of giant trees

Climate is one of the main drivers that delineates and influences biodiversity, geographic distribution, and the occurrence of plant species on a global scale. There is a latitudinal pattern especially in the tropics indicating that many plant species can be seriously affected by climate changes (Liang et al., 2022). However, there are still few approaches addressing why tropical species are more vulnerable to climate changes (Wiens et al., 2009). In the past, climatic conditions have significantly influenced the potential occurrence and formation of suitable areas for tall trees in the Amazon biome. For example, the pattern observed in the province of Roraima may reflect a combination of climatic and geological drivers acting directly over time, shaping the available ecological niches and the widespread distribution of tall trees. In particular, periods of more favorable climatic stability may have contributed to the dispersal and growth of giant tree species (Slik et al., 2024), while abrupt variations in climatic conditions have resulted in a more restricted mosaic of suitable areas. Additionally, low minimum temperatures during historical periods may have restricted the occurrence of tall tree species in certain regions (Larjavaara, 2014). Petrie et al. (2016) suggest that fluctuations in minimum temperatures affect processes such as regeneration and seedling establishment, which are critical phases for maintaining populations. Thus, areas with historically milder minimum temperatures would have been more favorable for the occurrence of many larger species.

These historical distribution patterns contrast with future climate scenarios, which indicate a reduction in the suitability of environmental conditions for giant trees and species. The climate variables analyzed, such as the maximum temperature of the hottest month, temperature seasonality, Isothermality, and precipitation regimes, play a crucial role in the development and survival of these trees in the projected scenarios. For example, increases in average temperatures and more significant variability in precipitation may result in a reduction in areas suitable for developing these trees, especially in the Xingu-Tapajós, Madeira, and Imerí provinces.

In the SSP1-2.6 scenario, which reflects a low-emissions future driven by global mitigation efforts to curb climate change (van Vuuren, Stehfest, et al., 2011), the projected impacts on species may be less severe. This is due to the ecological plasticity and adaptive traits of species, which could lead to habitat expansion into previously unsuitable areas. Notably, regions in the northern Amazon, such as the Guiana Shield, Roraima, and Pará provinces, are expected to present increasingly favorable conditions. Moderate temperatures, controlled seasonality, and adequate precipitation levels during the driest months support the fundamental physiological processes of these trees. This environmental stability allows for better thermal and hydric plasticity, enhancing growth rates and improving water use efficiency, thus enabling species to adjust effectively to changing climatic conditions (Smith et al., 2023). Isothermality, on the other hand, by remaining stable, ensures adequate thermal conditions throughout the year, which can avoid extreme stresses for giant trees, allowing greater photosynthetic efficiency and water transport (Becklin et al., 2021). Regular rainfall during the wettest quarter, particularly in regions such as the Province of Guyana and Roraima, ensures a continuous water supply for these trees, preserving the ecophysiological functions essential for their longevity and ability to regenerate.

The SSP5-8.5 scenario, which represents a future of high emissions and lack of control over climate change (Riahi et al., 2011; van Vuuren, Edmonds, et al., 2011), predicts a drastic reduction in habitat suitability for giant trees by 2080. The combination of extreme increases in the maximum temperature of the hottest month and increased temperature seasonality imposes severe thermal stress, which limits the resilience and plasticity of these trees to adapt to a hotter and drier environment (Allen et al., 2010; Olson et al., 2018; Sperry & Love, 2015). This scenario favors increased transpiration and difficulty in transporting water to the upper parts of the tree (Bartholomew et al., 2022; Liu et al., 2019), which can compromise their survival, especially in the most sensitive areas of the biome. Reduced precipitation in the driest month and during the hottest quarter in regions such as Rondônia and Xingu-Tapajós Province exacerbates water stress, which can result in higher mortality rates among giant trees due to decreased water availability and loss of ability to adapt to water variations (Stovall et al., 2019; Van Der Sande et al., 2023; Van Passel et al., 2022). The areas most severely affected by the loss of suitable habitat are located in the southern Amazon, where the effect of deforestation and fragmentation can also accelerate environmental degradation.

In the projected scenarios, a clear trend of reduction in habitat suitability is observed, particularly until the end of the 1980s in the SSP5-8.5 scenario. At the same time, in SSP1-2.6, some regions still maintain favorable conditions. The maximum temperature of the hottest month and precipitation during the driest month will also be determining factors that may limit the distribution of giant trees. The ability of these trees to cope with the projected extreme climate conditions will depend on the resilience of their physiological functions, such as transpiration, water transport, and photosynthesis, which are directly affected by temperature and water availability (Bennett et al., 2015; Kenzo et al., 2015). Analysis of these variables suggests that, without robust mitigation actions, the habitat of giant trees in the Amazon will be significantly reduced, especially in areas that already suffer anthropogenic pressures (Butt et al., 2023).

Although projections for tall trees show a worrying scenario, with a significant reduction in habitat suitability across much of the Amazon, analyses at the species level show important contrasts. *Dinizia excelsa*, for example, appears to find favorable areas in future scenarios, mainly in specific regions that may serve as climate refuges. *Goupia glabra* also maintains considerable suitability across large areas. These results show that the response to climate change varies among species, reflecting differences in environmental tolerance, current distribution, and potential for adaptation. For example, some areas of the Amazon may present conditions that favor species adapted to drier environments or with more significant seasonal variability (Brandão et al., 2022; Marengo et al., 2018; Nunes et al., 2022). Furthermore, a reasonable hypothesis is that with the reduction of other species that are less tolerant to climate change, *Dinizia excelsa* and *Goupia glabra* may occupy vacant niches, expanding their distribution into areas previously dominated by more sensitive species. Reproductive strategies, in particular, can be linked to the development of these larger species, as plants that are dispersed by wind can benefit from growing taller to maximize their geographical influence (Slik et al., 2024; Tuomisto et al., 2003). However, this potential habitat expansion reinforces the need to understand how specific ecological characteristics influence the resilience of Amazonian trees, providing support for conservation and forest management strategies in the face of climate change.

### 4.3 Limitations of the Study

A central limitation of this study concerns the methodological and conceptual differences between the datasets used in climate niche modeling. The discrepancy observed between habitat suitability projections for tall trees (height ⩾ 60 m), obtained via LiDAR, and large trees (dbh ⩾ 70) of *Dinizia excelsa* and *Goupia glabra*, derived from forest inventories and GBIF data may be related to distinct ecological, physiological, and structural factors that affect the distribution of these groups in different ways. While LiDAR data capture emergent trees regardless of species, occurrence data represent specific taxa, reflecting ecological adaptations and differentiated responses to climate change (Elith & Leathwick, 2009; Mäyrä et al., 2021).

Niche modeling based on tall trees via LiDAR may have underestimated the group’s resilience to climate change due to the broad variability of species involved. Since this dataset does not distinguish individual species, the resulting model represents an average ecological response of multiple emergent species, some of which may be highly vulnerable to environmental changes (Scheffer et al., 2018; Tiansawat et al., 2022; Wiens et al., 2009). In contrast, models based on the occurrence data of *Dinizia excelsa* and *Goupia glabra* capture the distribution of specific species whose ecological traits may confer more excellent stability to projected climate changes. This phenomenon suggests that these species’ ecological and physiological plasticity may allow them to persist under future climate conditions (Esquivel-Muelbert et al., 2019). At the same time, the emergent tree functional group as a whole may be subject to more significant environmental constraints.

Based on the geographic distribution of the data presented in the figure, it is evident that the occurrence points of tall trees mapped by LiDAR have broad coverage across the Amazon biome, spanning multiple biogeographic provinces. This extensive coverage suggests that climate niche modeling for emergent trees was not restricted to specific regions but was instead built on a dataset representative of the biome’s environmental heterogeneity (Gorgens et al., 2021). However, the occurrence data for *Dinizia excelsa* and *Goupia glabra*, obtained from forest inventories and GBIF, show distinct spatial patterns, with a higher concentration in certain provinces, such as the Guiana Shield and Pará (de Lima et al., 2022). This differentiated distribution may have resulted in a more stable niche model for these species, reflecting an ecological adaptation specific to the environmental conditions prevailing in the areas where they were recorded (Franklin, 2023). Thus, the sampling bias may not be in the extent of the LiDAR data but rather in the difference in spatial representativeness between the datasets, which may have influenced the habitat suitability projections in different ways for emergent trees and individual species.

Furthermore, LiDAR-based modeling focuses exclusively on the structural attributes of vegetation, whereas species occurrence data modeling directly captures the ecological preferences of these taxa (Coomes et al., 2017; Donager et al., 2018). Non-climatic factors, such as water table depth, soil fertility, and anthropogenic disturbances, can significantly influence the distribution of emergent trees but were not considered in the climate modeling. This aspect may have made the projections for emergent trees more sensitive to climate variations, as their occurrence may be linked to environmental factors not included in the models used.

Finally, the methodological approach to climate niche modeling contributed to the distinct results. Models trained with LiDAR data can capture spatial patterns of tree height but not necessarily the environmental conditions that favor specific species. In contrast, models based on species occurrence data are directly calibrated with environmental variables associated with the presence of individuals, which may result in more robust predictions for specific species, even under climate change scenarios.

### 4.4 Conservation implications and concluding observations

The tallest trees in the Amazon play crucial ecological roles, such as storing large amounts of carbon, regulating the microclimate, and maintaining biodiversity (Birdsey et al., 2023; Lindenmayer & Laurance, 2016; Pinho et al., 2020). Despite their importance, they are particularly vulnerable to changes in climate conditions. As a result, the sensitivity of these trees to climatic factors, such as Isothermality, maximum temperature of warmest month, and temperature seasonality, suggests that changes in these parameters can significantly alter their distribution and viability. For example, giant trees have a highly efficient vascular system, making them vulnerable to sudden temperature fluctuations and drought (Esquivel-Muelbert et al., 2019). Extreme temperatures and prolonged droughts can cause xylem embolism, leading to the collapse of water and nutrient transport, directly impacting their survival (Araújo et al., 2024; Bennett et al., 2015; Millard & Grelet, 2010).

In addition, the increase in natural disturbances associated with climate change can significantly impact the occurrence and survival of giant trees. That is the frequency of anomalous events, such as storms with strong winds and increased incidence of lightning, has already been observed (Gora et al., 2020; Gorgens et al., 2021) and has direct negative impacts on the tall trees. Therefore, analyzing the climatic variables associated with these trees’ occurrence and the projections generated by the climate niche model provides valuable information for the conservation and sustainable management of these species in the face of global changes. Overall, our analyses confirmed a clear spatial trend in the potential distribution of giant trees, indicating that the balanced combination of moderate daily temperature variations and minimum water availability during dry periods strongly influences suitable habitats for the tall trees, *Dinizia excelsa* and, Goupia glabra. In the projected climate change scenarios, which contemplate both optimistic and pessimistic scenarios, suitable habitats for these trees will be considerably reduced over the coming decades. In pessimistic scenarios, especially in the projection for 2061-2080, there is an apparent contraction of suitable areas, with regions of the provinces of Roraima and the Guiana Shield, where there is currently an abundance of these trees, being particularly affected. This suggests that these sites may become critical refuges for conserving these trees. Under the most pessimistic scenario, there is also a tendency for habitat suitability to shift to areas further north and west of the Amazon basin, implying habitat fragmentation (Brinck et al., 2017; Ma et al., 2023). This could lead to more significant pressure on the remaining populations, hindering their regeneration and amplifying the effects of the loss of ecological connectivity (Zurell et al., 2024).

Efforts to analyze the elasticity and plasticity of giant trees in more detail are needed in many areas of the biome. This information could provide more refined answers about how these trees respond to climate variations and their adaptive ways of adjusting their ecophysiological functions to maintain growth and reproduction. Thus, both the underlying and functional mechanisms are essential to predict the resilience of these trees in the face of environmental changes and to guide conservation and forest management policies in the Amazon. Understanding how past climatic conditions shaped the ranges of these trees is crucial for planning conservation strategies in the Amazon. Incorporating this information is essential to ensure the long-term survival of many giant species.

In summary, under the current scenarios projected by the sixth climate report of the Intergovernmental Panel on Climate Change, the average global temperature is estimated to increase by 1.5 °C between 2030 and 2052 and, if it continues to increase, precipitation levels will decrease, and drought scenarios in the Amazon will unfortunately become more frequent, directly impacting all biodiversity (Flores et al., 2024). Even in this scenario, only approximately 15% of the Brazilian Amazon is protected by conservation units, covering only about 58% of the remaining vegetation (IBGE, 2019). Given the ecological importance of giant trees, understanding the effects of climate change on their occurrence patterns is essential to refine conservation prospects in a changing world—for example, to what extent are protected areas in the Amazon currently susceptible to impacts induced by climate disturbances or anomalies, and how can giant trees withstand or respond to these changes? Large-scale ecological restoration efforts and the adoption of efficient public policies to mitigate the impacts of the climate crisis are urgently needed in light of the high and increasing rates of deforestation that devastate the world’s largest tropical forest and threaten the sanctuaries of the planet’s tallest trees.

## Authors’ contributions

R.B.L. conceived the main idea of the manuscript and accessed relevant Brazilian Amazon plots with the occurrence of large trees database with the approval of data owners. R.B.L. and E.B.G led the compilation of field inventory and remote sensing data with the assistance of C.P.O., D.A.S. S., D.G.S., L.A.G., C.C.V., J.R.M.F., A.P.B.B., J.P.G.S., and I.J.C.L., J.C.A., P.S.A. All co-authors contributed data. R.B.L. performed the analyses with the assistance of E.B.G. R.B.L. wrote the first draft of the manuscript, with all authors providing editorial input.

## Acknowledgements

Funding was provided by the Coordenação de Aperfeiçoamento de Pessoal de Nível Superior Brasil (CAPES; Finance Code 001); Conselho Nacional de Desenvolvimento Científico e Tecnológico (Processes 444350/2024-1 and 301432/2022-8); Amazon Fund (grant 14.2.0929.1); Universidade Federal dos Vales do Jequitinhonha e Mucuri (UFVJM); Instituto Chico Mendes da Biodiversidade and Instituto Nacional de Pesquisas Espaciais (INPE); Universidade do Estado do Amapá (Processes 0022.0279.1202.0018/2021).

